# TRPC6 channel inhibition disturbs store-operated Ca^2+^ entry to delay proliferation in bladder cancer

**DOI:** 10.64898/2026.01.29.702689

**Authors:** Jinhui Zha, Xiaotong Guo, Wentao Liao, Peitao Wu, Yuhong Wu, Yingyue Guo, Li Gao

**Affiliations:** Department of Urology;The Second Affiliated Hospital of Guilin Medical University, Guilin, 541000, China; Key Laboratory of Tumor Immunology and Microenvironmental Regulation, Guilin Medical University, Guilin, Guangxi, 541199, China; Guangxi Health Commission Key Laboratory of Tumor Immunology and Receptor-Targeted Drug Basic Research, Guilin Medical University, Guilin, Guangxi, 541199,China; Department of Urology;Shenzhen Nanshan People’s Hospital, Affiliated Nanshan Hospital of Shenzhen University,Shen zhen, 518052, China

**Keywords:** Transient receptor potential canonical type 6, Bladder cancer, Store-operated calcium entry, PI3K/Akt pathway, Proliferation

## Abstract

**Background and Objective:** Despite its prevalence, bladder cancer (BC) remains an unsolved pathogenesis. It is believed that TRPC6 channels have unique electrophysiological properties that contribute to intracellular Ca2+ signalling and tumorigenesis in cells. However, the mechanism by which TRPC6 contributes to BC progression and intracellular Ca2+ homeostasis remains unclear.

**Method:** In this study, TRPC6 expressions in paired BC and adjacent normal tissues were measured by immunohistochemistry. A KEGG pathway enrichment analysis was conducted to determine TRPC6’s potential contribution to BC. Ca2+ imaging analysis was performed to explore the contribution of TRPC6 in the BC cell. Flow cytometry and Cell Counting Kit-8 assay were performed to explore the effects of TRPC6 on the proliferation of BC. The impacts of TRPC6 SOCE on PI3K/Akt/mTOR pathway were measured by western blot. Based on the above bunch of studies, TRPC6 was found to be overexpressed in human BC tissue, which correlated with poor survival rates for patient overall survival (OS). We used the TRPC6-specific antagonist SAR7334 to explore and reveal significant inhibition of BC cell proliferation. Mechanistically, TRPC6 mediated cytosolic Ca2+ and regulation of SOCE, leading to the activation of the IP3K/AKT/mTOR pathway.

**Results:** We found that SAR7334 arrested the phosphorylation of PI3K and Akt, thus causing a significant decrease in phosphorylated mTOR. Similar effects were observed for the SOCE-specific antagonist MRS1845. In contrast, the Akt inhibitor MK2206 did not alter the SOCE in BC cells.

**Conclusions:** our results indicate that the pharmacological inhibition of TRPC6 arrests tumour cell proliferation through SOCE targeting the PI3K/Akt/mTOR pathway.

**Graphical abstract:** 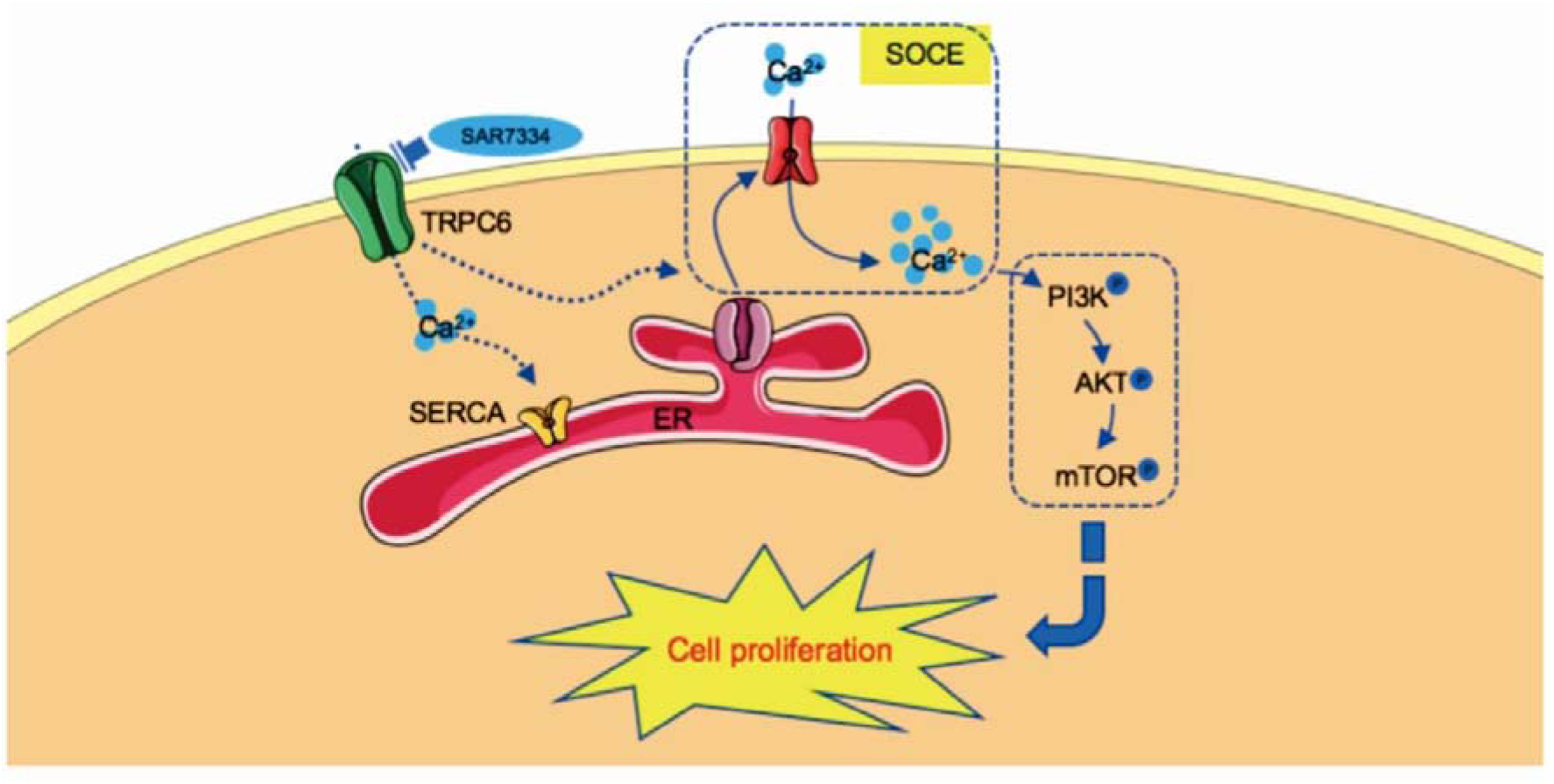

Schematic of proposed pathway. A model for summarizing TRPC6 contributes to cell proliferation in bladder cancer. TRPC6 specific inhibitor SAR7334 alter cellular cytosolic Ca2+ and SOCE, thus arrested phosphorylation of the PI3K/Akt pathway and the proliferation of bladder cancer cells.

## 1. Introduction

Bladder cancer (BC) is a large societal burden, with over 430,000 men and women diagnosed worldwide every year [1–3]. A key part of BC treatment is chemotherapy, especially for patients with non-muscle invasive BC. The use of intravenous chemotherapy after transurethral resection of BC can significantly reduce recurrence events and improve patient outcomes [4–6]. Even though chemotherapy and surgery have greatly improved the health of patients, the incidence and recurrence rates of BC remain high. It is important to explore the mechanism of tumorigenesis to BC and to find new anti-tumour drugs for the clinical treatment of BC.

The Ca^2+^ signalling pathway plays a major role in physiological and pathological cellular responses, such as gene expression, cell proliferation, migration, and invasion, which may contribute to cancer development [7, 8]. Intracellular Ca^2+^ is tightly regulated by influx/efflux mechanisms via ion channels, ATPase pumps, and exchangers, maintaining low intracellular Ca^2+^ levels to achieve homeostasis [9, 10]. In advanced cancer cells, altered Ca2+ flux leads to hypercalcemia and aberrant calcium signalling [11]. In these cases, transient receptor potential (TRP) channel is one of the pathways for calcium influx into cells, and channel activation leads to transient elevation in the concentration of intracellular Ca^2+^ [12]. It has been shown in numerous studies that TRP channels, especially those in the TRPC, TRPM, and TRPV subfamilies, are critical for tumour progression. Among these, TRPC6 is significantly associated with the progression of breast, gastric, oesophageal, and bladder cancers, as well as melanoma [13–18]. In this way, TRPC6-specific drugs may contribute to cancer treatment, given the critical role of TRPC6 in cancer-related cell signalling pathways, including proliferation, differentiation, and apoptosis [19–21]. Recent studies have shown that cancer vulnerabilities can be exploited not only by directly targeting constitutively expressed oncogenic drivers, but also by inducing pathway dependencies through epigenetic or signaling reprogramming [22]

On the other hand, in non-excitable cells, store-operated Ca^2+^ entry (SOCE) mediated by ORAI and the stromal interaction molecule (STIM) constitutes an important Ca^2+^ influx pathway [23]. In response to Ca2+ depletion, the endoplasmic reticulum-resident Ca^2+^ sensor protein STIM is activated to gate and open the ORAI Ca^2+^ channels resulting in Ca^2+^ influx within a few seconds [24–26]. It has been found that the expression levels of STIM and/or ORAI are correlated with tumour progression and poor prognosis in breast, gastric, oesophageal, and prostate cancers [24]. Correspondingly, SOCE mediated by the STIM and ORAI proteins alter Ca^2+^ signalling, involving various processes during oncogenic transformation such as cell apoptosis, proliferation, angiogenesis, and metastasis [27, 28]. In recent years, there has been a growing body of evidences indicating the role of TRP channels in the conduction of SOCE, especially the TRPC family [7]. The strongest evidence for the contribution of TRPC channels to SOCE has been described for TRPC1, TRPC4, and TRPC6, while TRPC-mediated SOCE appears to be dependent on cell type and level of expression [29–33].

In this study, we hypothesised that TRPC6 channels regulate BC progression through Ca^2+^ signalling via SOCE, postoperative biopsies and cell lines of BC patients were evaluated to test this hypothesis. We also explored the role of TRPC6 in SOCE generation in BC cells. Moreover, we used pharmacologic tools *in vitro* to explore the contribution of TRPC6-SOCE to the potential target pathway PI3K/Akt/mTOR. The clarification of this hypothesis will be crucial to understanding the role of TRPC6 in BC and may provide new therapeutic avenues.

## 2. Materials and Methods

### 2.1 Tissues and Cell lines

We analyzed 63paraffin-fixed specimens of human bladder urothelial carcinoma tissues and 16 neighboring conventional bladder transitional cell tissues collected between May 2007 to November 2011 from Shanghai Outdo Biotech Co., Ltd. (Shanghai, China). Another 10 pairs of paraffin-fixed specimens of human bladder urothelial carcinoma tissues and neighboring conventional bladder transitional cell tissues collected between September to December in 2017 from patients hospitalized in the Qianhai Shekou Free Trade Zone Hospital. The adjacent normal control was collected at least 2 cm away from the tumor margin [34]. The collected cancer tissue and its adjacent normal tissue have been identified under the microscope. All patients with BC were diagnosed by histopathology and treated with radical cystectomy or tumor resection. Human BC cell lines (sw870, T24, 5637, and J82) were obtained from the Cancer Institute, Central South University and human normal urothelial cells (SV-HUC-1) were acquired from the department of life science, Hunan Normal University (Changsha, China). BC cell lines were cultivated in RPMI-1640 media (Gibco, USA) supplemented with 10% fetal bovine serum (Gibco, USA), 1% streptomycin, and penicillin, and the SV-HUC-1 cells were cultivated in F-12k media (Gibco, USA) supplemented with 10% fetal bovine serum (Gibco, USA), 1% streptomycin, and penicillin. All the cells were cultured under 5% atm CO_2_ at 37°C.

### 2.2 Immunohistochemical (IHC) staining and scoring analyses

Immunohistochemical staining was conducted as previously described [34,35]. Briefly, paraffin sections of bladder cancer tissues and normal tissues were first deparaffinized and hydrated. Microwave antigen retrieval was performed for all antibodies, and endogenous peroxidase activity was blocked by incubating the slides in 0.3% H_2_O_2_. After serial incubation with primary antibodies and secondary antibody, sections were developed with peroxidase and 3,30-diaminobenzidine tetrahydrochloride. The sections were then counterstained with haematoxylin and mounted in non-aqueous mounting medium. TRPC6 antibody (1:500; Cusabio Biotech, Wuhan, China) was used to detect TRPC6 expression in the specimens.

Two independent experienced observers blinded to the histopathology diagnosis and clinical information reviewed and rated the extent of formalin- and paraffin-fixed immunostaining. They had been exposed to professional training in immunohistochemical staining scoring. Scores were based on combination of the ratio of completely stained cancer cells to the staining power. Tumor cell ratio was graded as 0 for no positive tumor cells; 1, < 10%; 2, 10%–25%; 3, 26–75%; and 4, > 75% positive tumor cells. Meanwhile, the extent of staining was rated as 1 for negative staining; 2, weak staining (light yellow); 3, adequate staining (yellow brown); and 4, heavy staining (brown). The staining index was calculated by multiplying the fraction of positive tumor cells by the staining power score. The staining index (total 0, 1, 2, 3, 4, 6, 8, 9, and 12) was used to estimate TRPC6 expression in bladder tissues, with scores ≥ 6 and < 6 considered as high and low expression, respectively. Five areas ata magnification of 400× were chosen randomly, and each area was scored separately according to the staining index. The score of a paraffin slide was obtained by the average score of these five areas [34, 36].

### 2.3 Western blot analysis

Western blot was conducted as previously described [37,38]. Briefly, the cells were washed with phosphate-buffered saline (PBS) and then lysed using RIPA buffer (0.5% sodium deoxycholate, 0.1% SDS, 50 mM Tris, and 150 mM NaCl, pH 8.0) with protease inhibitor mixture (Roche, USA) and phosphatase inhibitors (Roche, USA) at freezing condition for 15 min. The protein levels were measured with BC Protein Assay Reagent kit (Thermo Scientific, USA). A 10% SDS-polyacrylamide gel was used to separate tissue lysate aliquots containing 20 μg protein. These were subsequently moved to PVDF membranes (Millipore), and the membranes were consequently blocked for 2 hours with TBST buffer with 5% skim milk at 22 °C, and incubated at 4°C with primary antibodies overnight. We then added peroxidase-conjugated secondary antibodies and performed ECL (Cell Signaling Technology,12757) visualization. Band enumeration was conducted using densitometric analysis software (Bio-Rad). GAPDH expression was used as the internal standard to standardize the expression of the supplementary proteins. The primary antibodies were as follows: anti-TRPC6 (1:500; Cusabio Biotech, Wuhan, China), anti-GAPDH (1:3000, Proteintech Group, Chicago, USA), anti-Actin (1:2000, Proteintech Group, Chicago, USA), anti-PI3K (1:5000, Proteintech Group, Chicago, USA), anti-p-PI3K (1:1000, Abcam, Cambridge, UK), anti-Akt (1:2000, Proteintech Group, Chicago, USA), anti-p-Akt (1:2000, Proteintech Group, Chicago, USA), anti-mTOR (1:1000, CST, USA), anti-p-mTOR (1:1000, CST, USA). The secondary antibodies were HRP-Goat-anti-Rabbit IgG and HRP-Goat-anti-Mouse Ig G (1:1000, Cell Signaling Technology, USA).

### 2.4 Functional analysis and co-expression of TRPC6 genes

The Gene Ontology (GO) and Kyoto Encyclopedia of Genes and Genomes (KEGG) pathway enrichment analyses were performed using the DAVID (Database for Annotation, Visualization, and Integrated Discovery) and the Gene Expression Proling Interactive Analysis (GEPIA) datasets. The Search Tool for the Retrieval of Interacting Genes/Proteins (STRING: v.11.0: https://string-db.org/) database was used to establish and perform the visualization of the protein-protein interaction (PPI) network, which evaluated the functional and physical relationships between TRPC6 proteins. Additionally, STRING database was used to predict the TRPC6-related molecular function in Homo sapiens, and the genetic interaction (GI) network between TRPC6 and SOCE-related genes were analyzed by GeneMANIA (http://genemania.org/).GO analysis included molecular function (MF), biological process (BP), and cellular component (CC).

### 2.5 Immunofluorescence

Immunofluorescence was conducted as previously described[39,40].T24 and J82 BC cells were grown on collagen-coated coverslips and incubated for 24 h. Cells were then fixed in 4% paraformaldehyde, permeabilized using 0.3% Triton X-100, and stained with V5 polyclonal antibody (Bio-Rad) diluted in 1% bovine serum albumin–PBS for 1 to 2 hat room temperature. Cells were washed repeatedly with PBS, incubated with anti-TRPC6 (4□, overnight, Cusabio Biotech, Wuhan, China), fluorescence-labeled secondary antibody anti-Rabbit-IgG (37□, 90 minutes, Proteintech Group, Chicago, USA), and DAPI (37□, 10 minutes, Wellbio, China).

### 2.6 Cytotoxicity and apoptosis assay

In cell viability assay, cells were seeded in 96-well plates at 2,000 cells per well. Once attached, they were prepared as per their individual study procedure. The cells were treated with SAR7334 for 24 h and 48 h. Subsequently, 20 µl Cell Counting Kit−8 reagent (CK04, Dojindo Kumamoto, Japan) was supplemented to individual wells and retained for 2 h. Vmax microplate spectrophotometer (Molecular Devices, Sunnyvale, CA, USA) was used to measure the absorbance at 450 nm wavelength. These assays were performed in triplicate.

For apoptosis assay, cultured cells were seeded and treated with SAR7334. The apoptosis assay was conducted as previous described [32]. Briefly, both floating and adherent cells were collected and washed with PBS. Cells were stained with annexin-V/FITC and propidium miodide for 15 min at room temperature in dark. The fluorescence was detected by flow cytometry with the acquisition criteria of 10,000 events for each sample, and the quadrants were set according to the population of viable, untreated samples. The data were analyzed using FACS Aria equipped with the Cell Quest Software (BD Biosciences)

### 2.7 Ca^2+^ imaging measurements

Ca^2+^ events were measured as previously described [41,42]. Cultured bladder cancer cells were placed onto a 96-well fluorescent plate reader and incubated with the Ca^2+^ indicators fluo-4 AM (1µM) for 60 min at room temperature in Ca^2+^-free Hanks’ solution. After loading, cells were washed with PBS solution for 10 min at room temperature. Cells were imaged in a bath solution containing (mM): 134 NaCl, 6 KCl, 1 MgCl_2_, 2 CaCl_2_, 10 glucose and 10 HEPES (pH 7.4, NaOH). Cells were imaged in a bath solution containing (mM): 134 NaCl, 6 KCl, 1 MgCl_2_, 2 CaCl_2_, 10 glucose and 10 HEPES (pH 7.4, NaOH). Preparation was illuminated at 490 nm, and fluorescence emissions above 515 nm were captured through a barrier filter. Ca^2+^ events analyses were performed off-line and maximal amplitude of Ca^2+^ fluorescence was normalized by the initial fluorescence value (*F/F_0_*).

### 2.8 Statistical analysis

The obtained data were resultant of at least 3 separate assessments and were presented as mean ± S.D. Significance was tested using ANOVA, between-group comparisons were performed using Student’s t-test. Survival arcs were drawn according to Kaplan-Meier analysis and evaluated using log-rank test. All statistical analyses were performed using SPSS 21.0 (SPSS Incorporated, Chicago) statistical software package, and P < 0.05 was considered statistically significant.

### 2.9 Ethical approval

This study is approved to be conducted in accordance with the submitted clinical research protocol and informed consent form, based on ethical principles from the following: “Ethical Review Measures for Biomedical Research Involving Humans” (2016), “Good Clinical Practice” (2020), “Guiding Principles for Ethical Review of Drug Clinical Trials” (2010), the WMA “Declaration of Helsinki”, and the CIOMS “International Ethical Guidelines for Health-related Research Involving Humans”,and were approved by the Medical Ethics Committee of Shenzhen Qianhai Shekou Free Trade Zone Hospital.The approval number is 2023KY-004-01K.

## 3. Results

### 3.1 High expression of TRPC6 in human BC

The expression of TRPC6 was examined in the postoperative biopsies of patients with BC using paired histologically normal bladder urothelial biopsies as a control with the aim of exploring the relationship between TRPC6 expression and BC.

Histological diagnoses were established based on the examination of haematoxylin- and eosin-stained specimens. Samples from bladder tumours showed higher expression of TRPC6 compared to the histologically normal controls via immunohistochemistry (Figure. 1A–B). The Kaplan–Meier survival curve showed that the overall survival of patients with high TRPC6 expression was lower than that of patients with low TRPC6 expression (Figure. 1C-D; P < 0.001). These results support the idea that upregulated TRPC6 expression is related to poor prognosis in BC.

**Figure 1.**
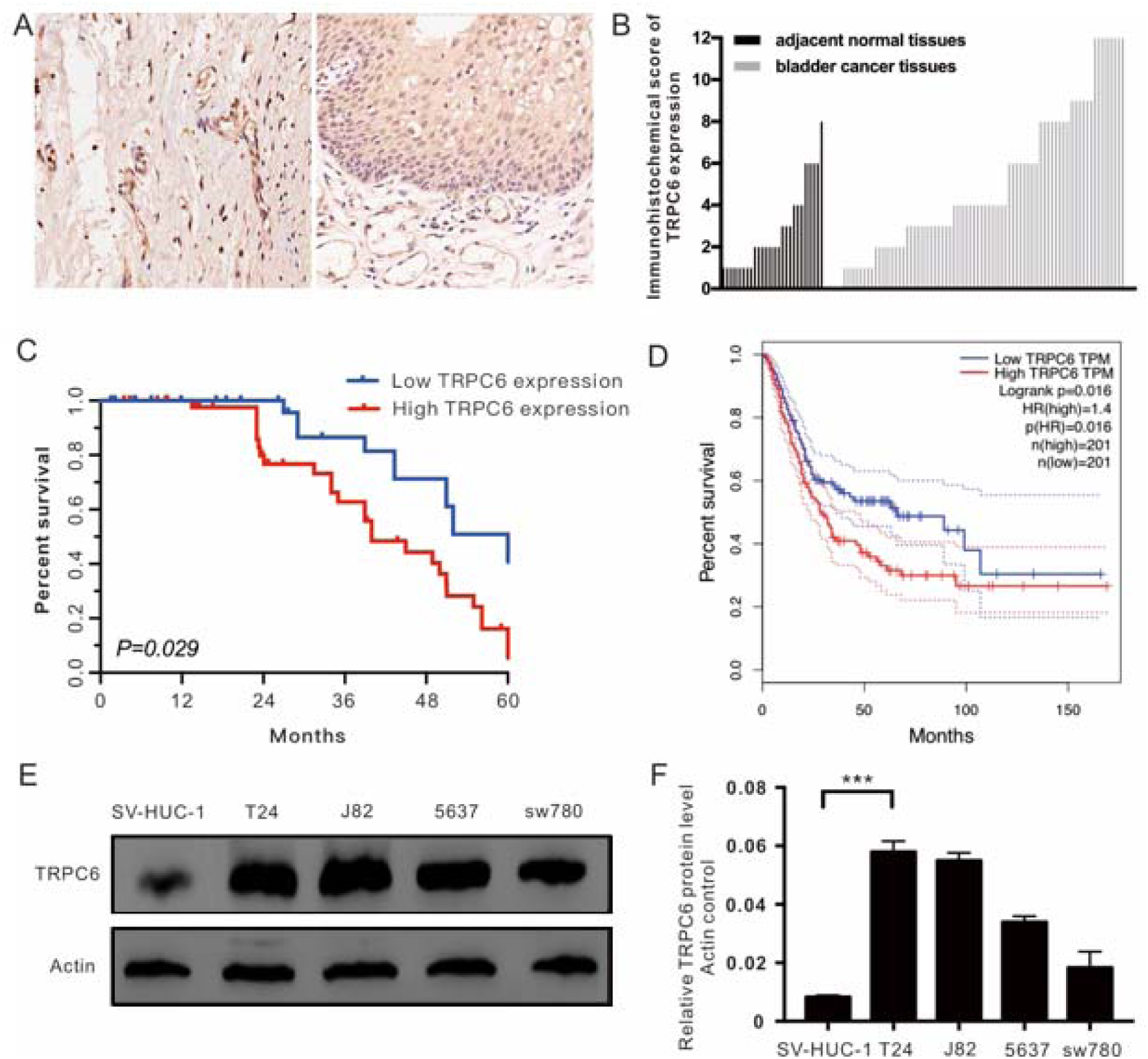
Expression of TRPC6 protein in BC. **A.** Expression of TRPC6 protein in BC tissues and adjacent normal bladder biopsies. **B.** Immunoreactive scores of TRPC6 relative changes. **C.** Overall survival in bladder cancer patients in our cohort. Patients were stratified according to the mean value of each measure in this population. **D.** is the same as **C.** but the data extracted from the GEPIA dataset. **E.** Western blot analysis of the levels of TRPC6 in the normal bladder urothelial cell line, SV-HUC-1, and different types of bladder cancer cell lines, T24, J82, 5637, and SW780.Original blots are presented in Supplementary Figure 1. **F.** Quantitative analysis of TRPC6 relative expression. ***Compared to the control; p<0.01.

We also compared TRPC6 expression in a normal bladder urothelial cell line, SV-HUC-1, and different types of bladder cancer cell lines—T24, J82, 5637, and sw780—using a western blot technique. Consistent with the immunohistochemistry findings in patients with BC, TRPC6 was higher in the T24 and J82 cells than in the cells derived from normal bladder tissues, although TRPC6 expression was more variable in 5637 and SW780 cells (Figure. 1E–F). Since T24 and J82 cells more closely reflect the overexpression of TRPC6 observed in human BC samples, these two cell lines were used for functional experiments in this study.

### 3.2 TRPC6□related gene-enrichment analysis

The protein-protein interaction (PPI) networks of TRPC6 were constructed, in which 51 nodes correlating with TRPC6 (Figure.2A) formed the networks with 383 edges (PPI enrichment p values:< 1.0 × 10−16; local clustering coefficient: 0.711). Meanwhile, the enrichment of molecular functions as determined through Gene Ontology (GO) analysis showed that TRPC6 significantly contributes to “calcium channel activity” (GO:0005262) and “store-operated calcium channel activity” (GO:0015279). Thus, we used GeneMANIA to establish a Genetic Interaction (GI) network, which revealed that there were strong physical interactions between TRPC6 and SOCE-related genes (Figure. 2B).

**Figure 2.**
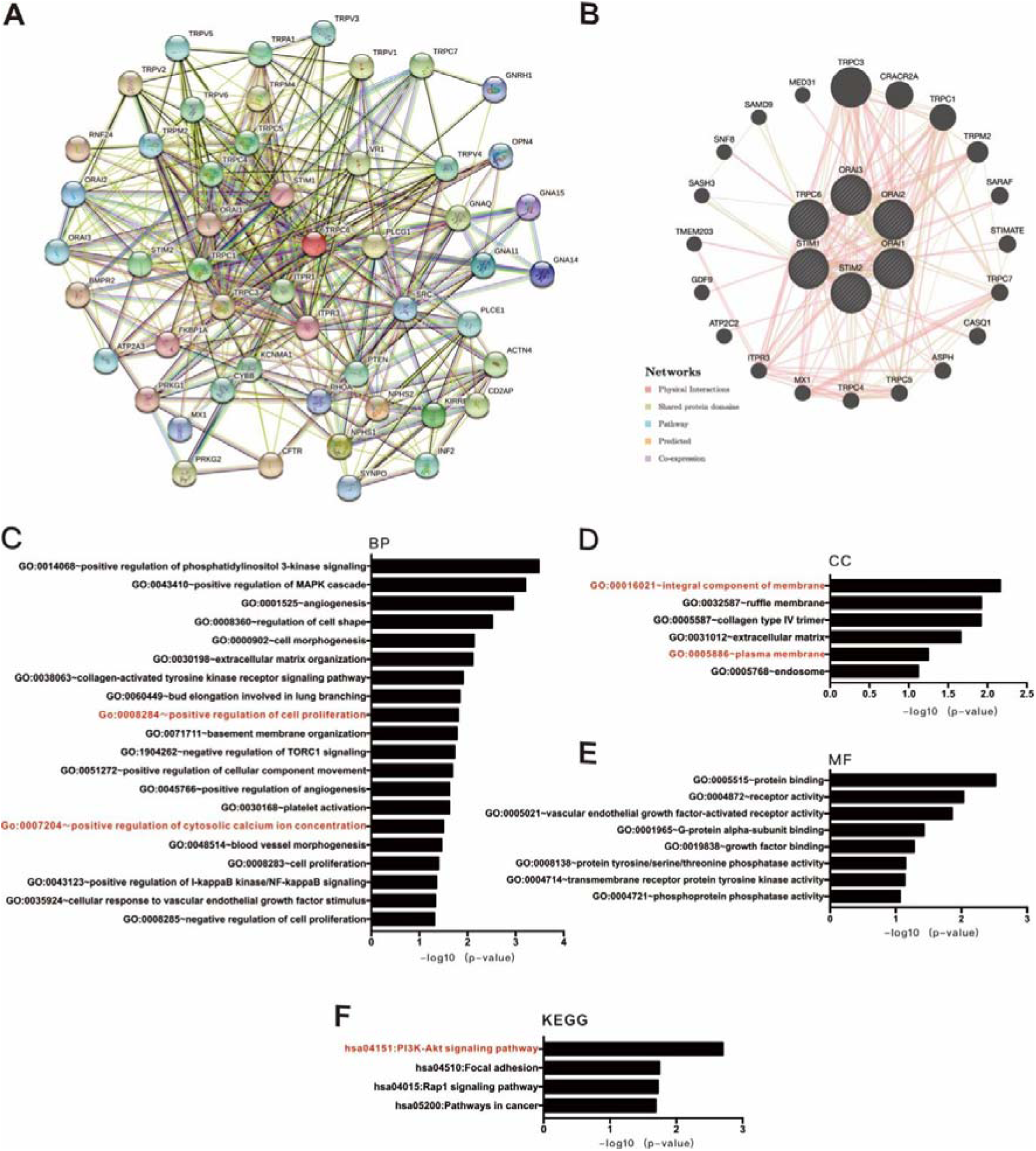
Functional enrichment analysis based on the TCGA dataset. **A.** Overview of the PPI network constructed using STRING 10.0 database. The network includes 51 nodes correlating with TRPC6 (Fig. 1A), which formed the networks with 383 edges (PPI enrichment p values:< 1.0 × 10−16; local clustering coefficient: 0.711). **B.** GI network for TRPC6 and SOCE-related genes established by GeneMANIA. **C–F.** KEGG and GO analyses for TRPC6 genes in BC using DAVID tools. (PPI, protein-protein interaction; GI, genetic interaction; SOCE, store-operated calcium entry; KEGG, Kyoto Encyclopedia of Genes and Genomes; DAVID, database for annotation, visualisation, and integrated discovery; GO, gene ontology; BP, biological process; CC, cellular component; MF, molecular function).

The top 50 genes that were positively correlated with TRPC6 in BC tissues (upregulated DEGs; uDEGs) were determined using the TCGA dataset and the GEPIA web-based tool. The uDEGs were inputted into the DAVID, where the KEGG pathways and GO function were analysed to identify the biological functions of the TRPC6 genes (Figure. 2C–F). KEGG BRITE functional hierarchies for TRPC6 showed that the co-upregulated genes had a preponderance of genes representing the ‘PI3K-Akt signalling pathway’, ‘Focal adhesion’, ‘Rap1 signalling pathway’, and ‘Pathways in cancer’, among others (Figure. 2F).

GO function enrichment was performed on the genes that co-expressed TRPC6; the DAVID tool was used to analyse their possible activities in biological processes, molecular functions, and cellular components. Based on the GO analysis for TRPC6 and its co-expressed genes, the most enriched ontology terms as show in Figure 2C-D.

### 3.3 Plasma membrane position TRPC6 contributes to Ca^2+^entry in BC

TRPC6 is a non-selective channel that mainly conducts Ca^2+^ and mediates cytosolic Ca^2+^ concentrations. These processes play a major part in cell-cycle progression, cell proliferation, and cell division. The above results indicated that TRPC6 was involved with cytosolic Ca^2+^regulation (GO:0007204∼positive regulation of cytosolic calcium ion concentration; Figure.2C). We here first tested the localization of TRCP6 in BC cells. By confocal microscopy, we showed that TRPC6 protein was expressed in the cytoplasm and plasma membrane of the T24 and J28 cells. We next studied how TRPC6 works in Ca^2+^ signalling in BC cells. We used the TRPC6-specific antagonist SAR7334 to evaluate TRPC6-dependent calcium influx in physiological condition (Ca^2+^, 2 mM). SAR7334 reduced the thapsigargin-evoked calcium efflux in a dose- and time-dependent manner (Figure. 3B–E). To obtain further evidence that the observed effects were due to decreased [Ca^2+^]_ER_ store, the experiments were repeated by using SAR7334 with a Ca^2+^-free medium. SAR7334decreased the thapsigargin-evoked calcium efflux in cytosolic Ca^2+^ concentration (Figure. 3E–F). These results indicate that TRPC6 is expressed in the plasma membrane and contributes to the cytosolic Ca^2+^concentration.

**Figure 3.**
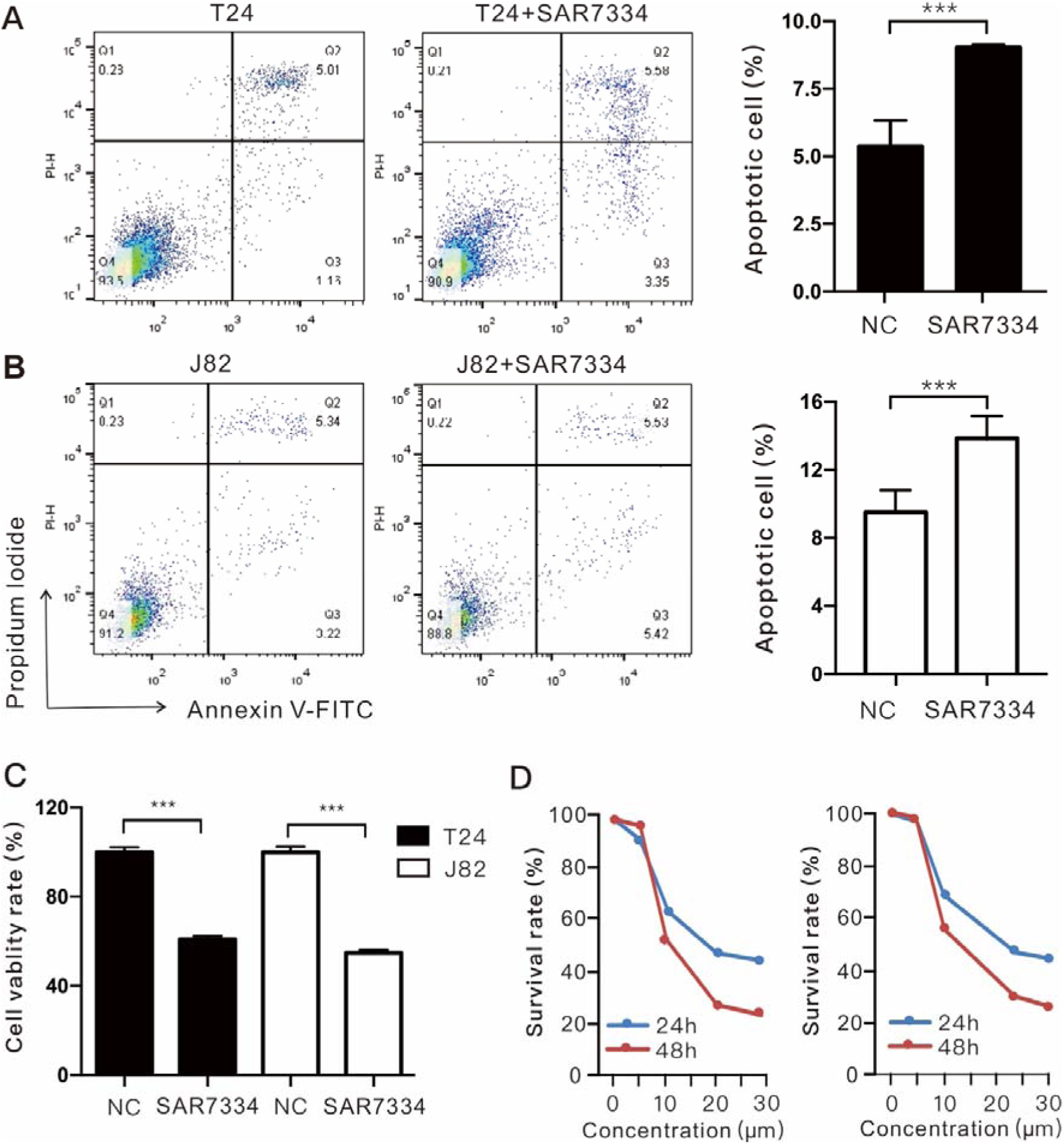
Plasma membrane localisation of TRPC6 and Ca^2+^ signalling in BC cells. **A.** TRPC6 is expressed and located in the cytoplasm and plasma membrane. **B–E.** Fura-4-AM fluorescence ratio of cells in response to thapsigargin (1 μM) stimulation and summary of the results. n =18–20 cells; 6–8 cells recorded and analysed from each group and repeated 3 times. SAR, SAR7334. *\*\*P < 0.01*.

### 3.4 Role of TRPC6 channels in SOCE in BC cell lines

SOCE has been reported to play an important role in supporting several cancers, including breast, colon, gastric, and oral cancers. As our results indicated that the TPRC6 channels blocked significantly attenuate thapsigargin-evoked calcium efflux, we hypothesized that TRPC6 might be playing a role in the conduction of SOCE regulating cellular cytosolic Ca^2+^ generation and resulting relevant features of cancer cells, such as proliferation and migration. We evaluated the contribution of TRPC6 in SOCE generation in BC cell lines using SAR7334. The cells were suspended in a Ca^2+^-free medium, and we found that thapsigargin-evoked calcium release was higher in the control cells compared to the cells treated with SAR7334. The subsequent addition of CaCl_2_ (2 mM) to the extracellular medium following thapsigargin-treatment resulted in a further cytosolic Ca^2+^ increase indicative of SOCE (Figure. 4).

**Figure 4.**
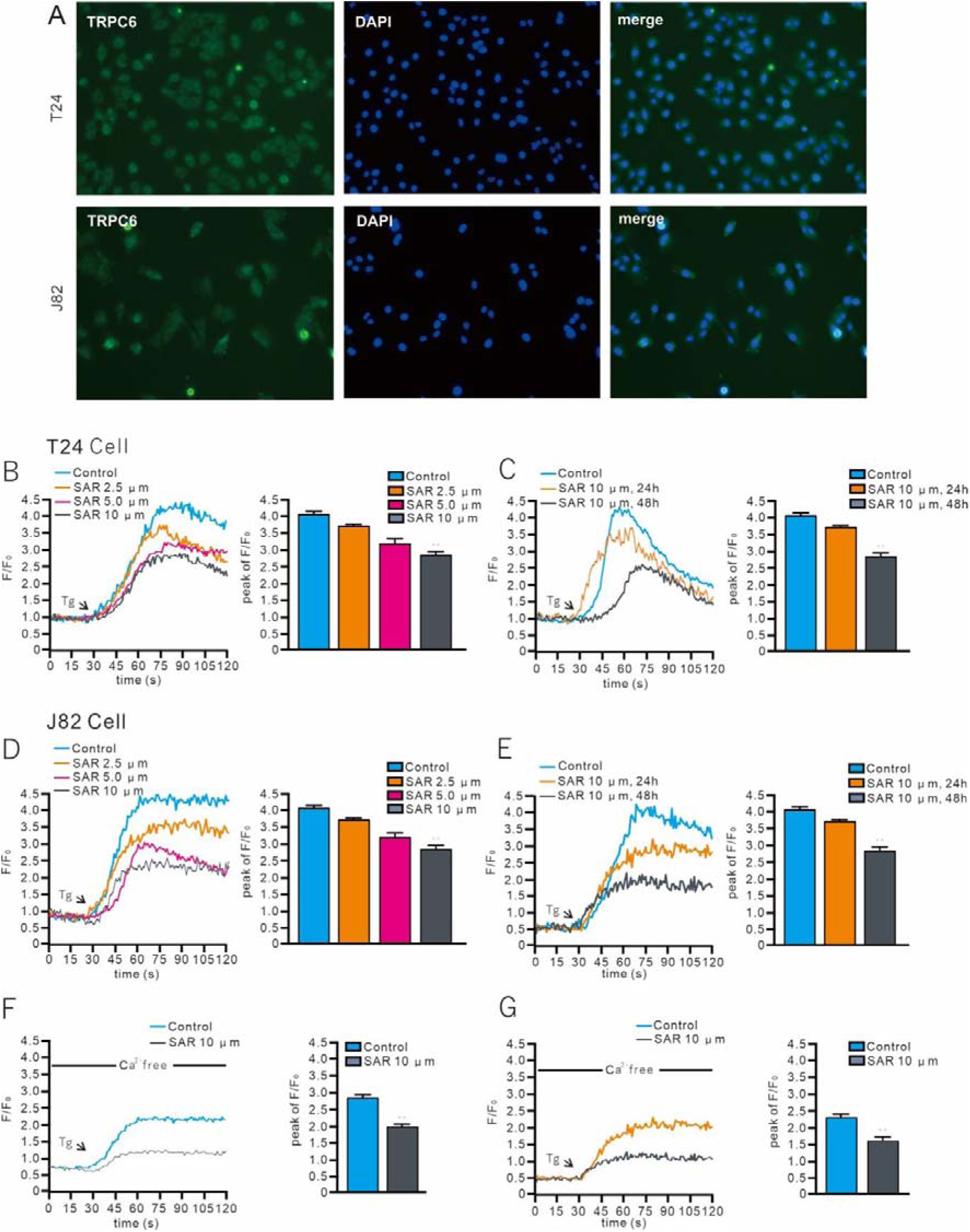
SAR7334 attenuates store-operatedCa^2+^ entry in BC cells. **A.** Ca^2+^ fluorescence images of a Fluo-4-AM–loaded T24 cell. **B.** Time course of Ca^2+^ fluorescence changes in the cellular ROI and summary of the results. Cells were perfused with a Ca^2+^-free medium and then stimulated with 1 μM thapsigargin followed by reintroduction of external Ca^2+^ (final concentration 2 mM) to initiate Ca^2+^ entry. **C.** Same as A, but in a J82 cell. **D.** Same as B, but extra from C. n=18-20 cells; 6–8 cells recorded and analysed from each group and repeated 3 times. SAR, SAR7334. *\*\*\*P < 0.001*.

### 3.5 Pharmacological inhibition of TRPC6 on BC cell proliferation

In the GO analysis, GO:0008284 (positive regulation of cell proliferation) was greatly related to the functions of TRPC6 in BC (Figure. 2C); therefore, SAR7334 was applied to test the contribution of TRPC6 on cell proliferation in the T24 and J82 cells. Flow cytometry and the trypan blue exclusion test showed that SAR7334 treatment significantly decreased the cell viability in the T24 and J82 cells (Figure. 5A–C). CCK8 assay showed that SAR7334 significantly slowed the growth speed of the T24 and J82 cells (Figure. 5D). These findings show that TRPC6 could stimulate the proliferation of BC cells.

**Figure 5.**
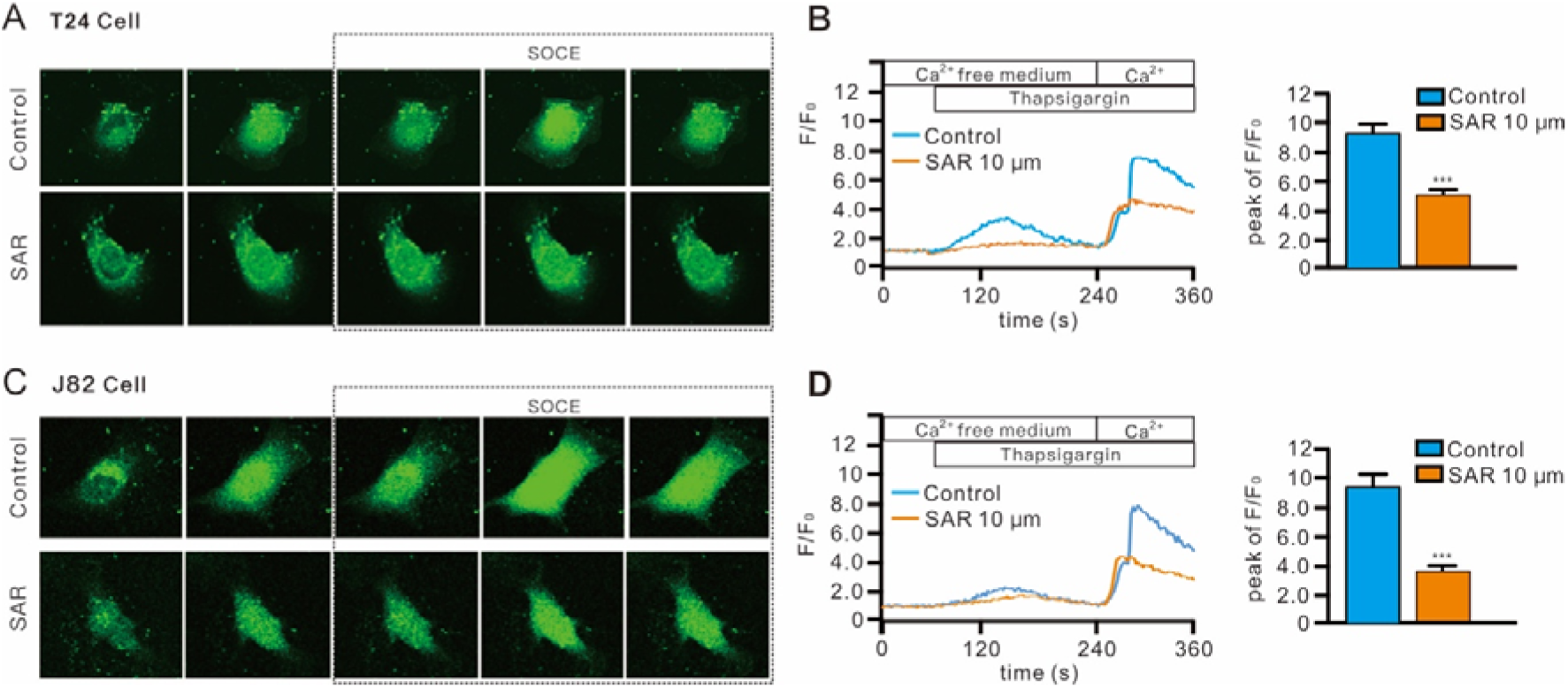
SAR7334 impairs the proliferation of BC cell lines. **A–B.** Representative images of flow cytometric analysis were used to show viability rates in cells treated with 10 μM SAR7334. **C.** Cell viability was determined by the trypan blue exclusion assay. **D.** CCK8 assays revealed the proliferation of indicated bladder cancer cells. The results are presented as the mean ± SD. *\*\*\*P < 0.001*.

### 3.6 TRPC6 regulates PI3K/Akt phosphorylation in BC cells

We explored the possible mechanism underlying the functional role of TRPC6-related SOCE in BC cells. In the KEGG analysis, hsa04151 (PI3K-Akt signalling pathway) was shown to be greatly related to the functions of TRPC6 in BC (Figure. 2F); thus, we examined the effects of TRPC6 on PI3K and Akt in BC. Western blots indicated that SAR7334 treatment arrested the phosphorylation of PI3K and Akt, thus subsequently causing a significant decrease in phosphorylated mTOR (p-mTOR). SAR7334 treatment, however, did not alter the total PI3K, Akt, and mTOR expression (Figure. 6). These results indicated that treatment with SAR7334 resulted in the inactivation of the Akt/mTOR pathway. We also evaluated the effects of SOCE on PI3K/Akt/mTOR phosphorylation using the SOCE blocker MRS1845. In the presence of MRS1845, the phosphorylation of PI3K/Akt/mTOR was lower in the T24 and J82 cells. Meanwhile, there were no significant changes in total PI3K/Akt/mTOR expression (Figure. 6). Together, it is likely that TRPC6-mediatedSOCE, enabling the activation of the PI3K/Akt/mTOR pathway, promotes the propagation of BC cells.

**Figure 6.**
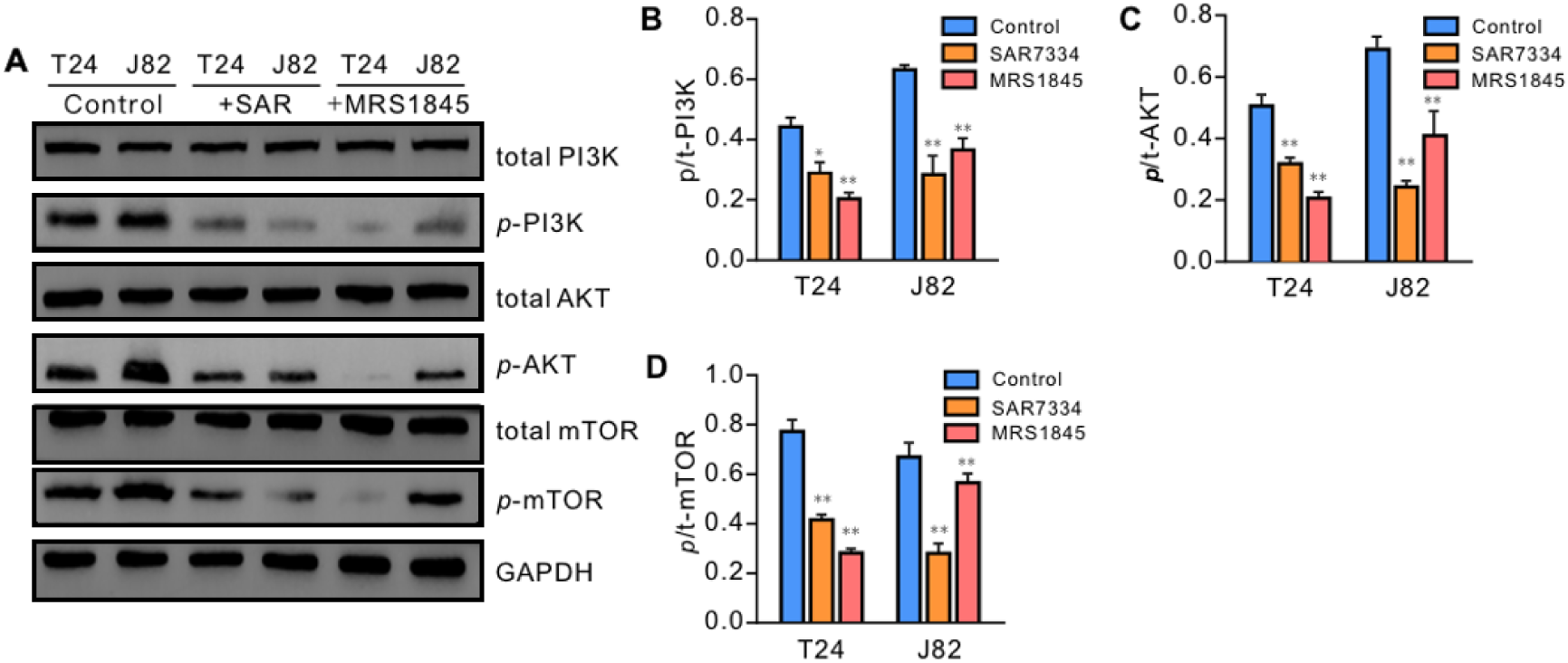
SAR7334-arrested phosphorylation of the PI3K/Akt pathway in BC cells. **A.** Representative western blots showing the alternation of the phosphorylated (p-PI3K, p-Akt, p-mTOR) components and total proteins (t-PI3K, t-Akt, and t-mTOR) of the PI3K/Akt signalling pathway induced by 10 μM SAR7334 and 1 μM MRS1845 stimulation. Protein expression was assessed at 48 hours after SAR7334 and MRS1845 treatment.Original blots are presented in Supplementary Figure 2. **B.** Summary data (means ± SE, n=3) were quantitated and compared, and the control was not treated byTRPC6.

### 3.7 Influence of AKT pathway on cell SOCE

To further confirm the relationship between SOCE and the Akt pathway in BC cells, we investigated the effect of inhibition on the PI3K/Akt/mTOR pathway with TRPC6-enhanced SOCE in T24 and J82 cells using the Akt-specific antagonist MK2206. We found that the MK2206 treatment did not alter SOCE that was induced by the passive depletion of the intracellular Ca^2+^ stores with thapsigargin, as compared to the control BC cells (Figure. 7A, C). In accordance with our former data, the treatment of BC cells with SAR7334 significantly attenuated the amplitude of SOCE. However, MK2206 did not further reduce the SOCE in SAR7334-treated BC cells.

**Figure 7.**
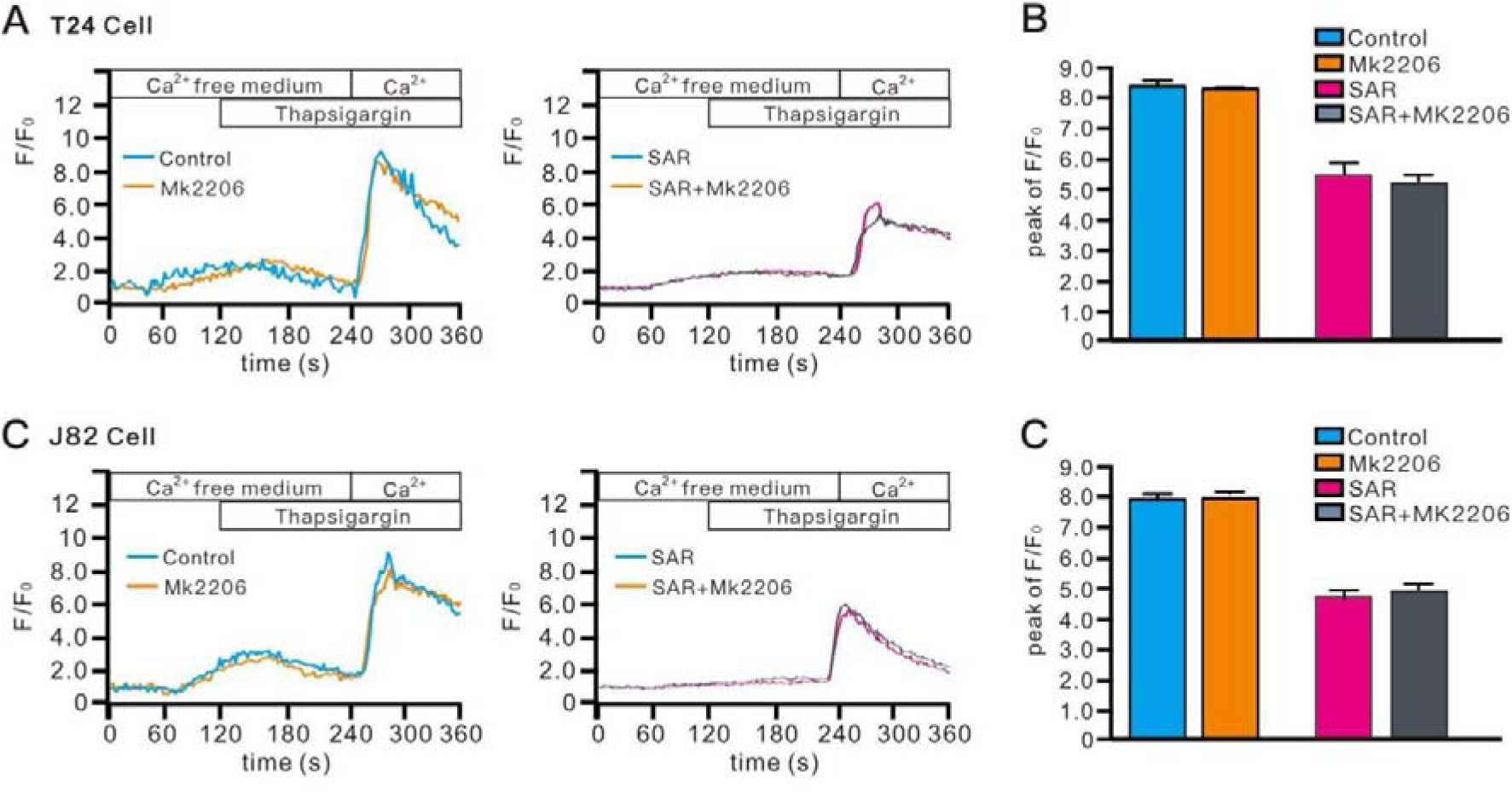
Inhibition of PI3K/Akt with a specific antagonist does not alter SOCE in BC. **A.** Representative traces showing changes in cytosolic Ca^2+^ concentration; expressed as the ratio of the Ca^2+^ fluorescence images of a Flo-4-AM-loaded BC cell) in response to thapsigargin (TG)-induced store depletion in the absence of extracellular Ca^2+^ and subsequent SOCE upon replenishment of extracellular Ca^2+^. Cells were treated with MK2206, SAR7334, and SAR7334+MK2206 for 48 hours before Ca^2+^ measurement, respectively.

## 4. Discussion

Our study found that as compared with normal tissues, human BC tissues expressed higher levels of TRPC6. We identified that the TRPC6 is involved in calcium signalling and regulates the generation of SOCEs in non-excitable BC cells. We used the TRPC6-specific antagonist SAR7334 to demonstrate that TRPC6 channels contribute to the proliferation of BC cells. This may be because TRPC6/SOCE mediates the phosphorylation of the PI3K/Akt pathway. Our study opens up the possibility that TRPC6 channels may be a potential target for BC therapy.

Ca^2+^-dependent signalling is important in cell proliferation, migration, and invasion [43–46]. Ca2+ homeostasis is frequently deregulated in cancer cells, and TRP channels may play a key role in this process [47]. TRP channel genes were abnormally upregulated in various types of tumours, suggesting their involvement in tumorigenesis [48–52]. TRPC6 channels are expressed in the human bladder, which belong to a family of non-selective cation channels [53]. The TRPC6 ion channels allow positive ions, especially calcium ions, into the cells, which activate a cascade of signalling events [53, 54]. For example, TRPC6-mediated Ca^2+^ signalling, which activates the calmodulin-dependent protein kinase and mitogen-activated protein kinase, is required for the proliferative response of cancer cells [55]. TRPC6 channels also play a role in SOCE, as both STIM/ORAI-regulated store-operated channels and as second messenger-operated channels [7, 56, 57]. Additionally, TRPC6 has been shown to play a key role in Ca2+-mediated proliferation of BC cells in recent study [15]. Nevertheless, the precise mechanisms by which TRPC6 contributes to BC progression in humans, as well as the mechanisms and precise dynamics of TRPC6’s regulation of Ca2+ homeostasis remain obscure. The TRPC6 protein expression in BC tissues was upregulated compared to normal bladder tissues, and TRPC6 expression was significantly associated with poor prognosis [15]. In cells, TRPC6 functions as a calcium channel and regulates cytosolic Ca2+ levels [7, 56, 57]. Our results indicate that TRPC6 inhibition drastically reduced thapsigargin-evoked Ca^2+^ entry into BC cells. Moreover, inhibition of TRPC6 in BC cells also resulted in a significant reduction in Ca2+-evoked Ca2+ entry, consistent with previous observations, which indicated that TRPC6-mediated Ca^2+^ signalling plays an important role in SOCE in these BC cell lines.

In non-excitable cells, SOCE is the main Ca2+ influx pathway, and is essential for cell function [58]. SOCE mediated by the STIM and ORAI proteins is involved in various cellular processes during tumorigenesis, such as malignant transformation, apoptosis, proliferation, angiogenesis, metastasis, and antitumour immunity [28, 59]. It has been reported that TRPC6 controls SOCE generation by regulating ORAI channels rather than by regulating Ca2+ entry, as in breast cancer [7, 16]. The same finding was observed in prostate cancer cells where TRPC6 expression was silenced in calcium-stimulated PC3 cells, resulting in a decrease in SOCE and proliferation [60, 61]. This study confirms previous findings, focusing on the cellular pathway by which TRPC6/SOCE promotes cell proliferation through the TRPC6/SOCE pathway, we were able to determine that TRPC6 regulates cytosolic Ca^2+^ and controls SOCE in BC cells. The mechanisms by which SOCE regulates the proliferation of cancer cells remain poorly defined. The proposed mechanisms include the regulation of the dephosphorylates Cdc2 and ERK or PKB/Akt signalling [62–64]. A KEGG pathway enrichment analysis revealed TRPC6/SOCE to be significantly correlated with the PI3K/Akt pathway in BC, revealing their importance in cell proliferation.Beyond aberrant expression, genetic variations such as single-nucleotide polymorphisms (SNPs) in ion channels and Ca²□ signaling genes have been linked to cancer susceptibility, progression, and treatment response [65–67].

In addition to its role in normal cellular functions, the PI3K/Akt pathway contributes to tumorigenesis as well, including cellular proliferation, growth, survival, and mobility [68, 69]. Recently, TRPC6 has been shown to promote cell apoptosis by stimulating the PI3K/Akt pathway via cytoprotective autophagy [70,71]. In this study, we showed that a TRPC6/SOCE mediates the proliferative effect of PI3K/AKT/mTOR on human BC cells. Initially, KEGG pathway enrichment analysis showed TRPC6-related genes are associated with the PI3K-Akt signalling pathway in BC. We demonstrated that the activity of the PI3K/AKT pathway is impaired in TRPC6 inhibition, as evidenced by the fact that the phosphorylation of the PI3K, AKT, and mTOR proteins was greatly reduced using a TRPC6 antagonist. The phosphorylation of Akt and p70S6K in mice lacking TRPC6 is consistent with recent reports [71]. In addition, It has been reported that the TRPC6 antagonist oleocanthal arrests the phosphorylation of Akt [72, 73]. The mechanisms underlying the crosstalk between the TRPC6/SOCE and PI3K-AKT-mTOR pathways in BC cells remain to be elucidated. Furthermore, The pharmacologic distortion of SOCE reduced PI3K/AKT/mTOR phosphorylation, whereas inhibition of AKT in BC cells did not alter SOCE generation. This was consistent with recent reports that SOCE is critical to the activation of the PI3K-AKT kinase-mTOR pathway, facilitating cell growth, cell cycle entry, and T-cell proliferation [74]. Moreover, PI3K/AKT phosphorylation is controlled by TRPC6 via Ca2+ signalling, suggestive of the role TRPC6 might play in signalling SOCE.

## 5. Conclusions

Our work shows that the expression of TRPC6 was greatly increased in human BC tissues and cell lines, which was correlated with poor patient prognosis. The pharmacological inhibition of TRPC6 induced decreased Ca^2+^ influx and SCOE generation in BC cells, which might be associated with reduced cell proliferation. Mechanically, TRPC6-regulated cytosolic Ca^2+^ and SOCE are critical in IP3K/Akt phosphorylation, which may contribute to the proliferation of human BC cells. These findings provide evidence that TRPC6 expression can be a predictor of poor prognosis in patients with BC and should be studied as a therapeutic target for novel molecular therapy.

## 7. Declarations

### Consent for publication

Not applicable.

### Data availability statement

All datasets generated for this study are included in the article/ supplementary material.

### Ethics statement

The studies involving human participants were reviewed and approved by The study involving human participants was approved by he ethics committee of the Qianhai Shekou Free Trade Zone Hospital, Ethical Batch Number:2023KY-004-01K. Patients/participants provided written informed consent for their participation in this study. The patients/participants provided their written informed consent to participate in this study.

### Author contributions

Weifeng Yang was involved in funding acquisition and research supervision. Jinhui Zha and Xiaotong Guo were responsible for sample collection and experiment execution. Yuhong Wu and Yingyue Guo carried out data analysis and visualization. Wentao Liao and Peitao Wu drafted the manuscript. Li Gao contributed to critical revision of the article. All authors have approved the final version of the manuscript and agree to be accountable for all aspects of the work. All persons designated as authors qualify for authorship, and all those who qualify for authorship are listed.

### Author Statements

All authors confirm their participation in this research and the development of the manuscript. Furthermore, all authors have reviewed and approved the final version of the manuscript and consent to its submission for publication.

